# Dysregulated DnaB unwinding induces replisome decoupling and daughter strand gaps that are countered by RecA polymerization

**DOI:** 10.1101/2023.06.26.546485

**Authors:** Megan S. Behrmann, Himasha M. Perera, Malisha U. Welikala, Jacquelynn E. Matthews, Michael A. Trakselis

**Author notes:** To whom correspondence should be addressed: *Michael A. Trakselis, One Bear Place #97348, Waco, TX 76798-7348.

## Abstract

Repair of DNA damage begins with the elicitation of targeted cellular responses to restore the genome. In *E. coli*, major products of DNA damage result in the buildup of single-stranded DNA (ssDNA) that is rapidly bound by cooperative filamentation of RecA to initiate the SOS response. The replicative helicase, DnaB, is a central component of the replisome, unwinding duplex DNA in concert with Pol III template dependent synthesis. Interestingly, helicase unwinding is heavily regulated, and the unwinding rate can be reduced by over 10-fold if DnaB becomes decoupled from Pol III. However, if DnaB is dysregulated by mutations that enforce a faster more constricted conformation, unwinding can continue independently, generating excess ssDNA resulting in severe cellular stress. This surplus ssDNA can stimulate RecA recruitment for recombinational repair or activation of SOS to increase the available repair protein pool. To better understand the consequences of dysregulated unwinding, we combined targeted *dnaB* mutations with an inducible plasmid-based RecA filament inhibition strategy to examine the dependencies on RecA in counteracting decoupling. We find that RecA filamentation is instrumental for processing daughter strand gaps left behind from decoupled unwinding and synthesis to prevent DNA breaks. Without functional RecA filaments, *dnaB* mutant strains had a greater burden from endogenous damage but without a compensatory increase in mutagenesis. Overall, RecA plays a critical role in strain survival by processing DNA gaps and protecting from breaks caused by dysregulated or interrupted helicase activity *in vivo*.

**AUTHOR SUMMARY:** Coupled DNA unwinding and synthesis is a genomic protection strategy used during DNA replication to prevent excessive buildup of labile single-stranded DNA (ssDNA). The helicase and polymerase enzymes have evolved multidimensional regulation tactics to maintain this connection despite different individual kinetic rates and differential responses to genomic obstacles. For one, the DnaB helicase in *E. coli* can alter its hexameric ring structure by dilating to slow down or constricting to speed up DNA unwinding. Here, we have utilized persistently constricted mutants of DnaB *in vitro* or genomically edited *dnaB in vivo* to induce decoupling in the replisome. Constricted DnaB mutants limit total leading strand synthesis by Pol III, indicating that lost kinetic regulation between these enzymes results in inefficient replication. Using an inducible plasmid-based system to disrupt Rad51 filamentation i*n vivo*, we show that RecA is responsible for the increased mutagenesis, a filamented cellular phenotype, and mitigating DNA breaks from excess ssDNA caused by decoupling. These results reveal a role for RecA filamentation in mediating excess ssDNA resulting from decoupling to maintain survival and adaptation.

## INTRODUCTION

Replicative hexameric helicase enzymes are at the forefront of DNA replication, a highly conserved process fundamental to all life. Yet, the mechanisms for how these enzymes regulate their activities and function in concert with other replisome proteins are not fully understood at a mechanical or biological level. While the individual roles of replication proteins, including helicases, are well known, the mechanisms with which they regulate interactions with DNA and other proteins in a cellular environment remain to be fully elucidated. The bacterial helicase, DnaB, is responsible for exposing single stranded DNA (ssDNA) strands as it creates the “fork” around which replisome proteins orient [1, 2]. DnaB is a RecA-like superfamily 4 (SF4) helicase, translocating and unwinding in the 5’-3’ direction on the lagging strand with a C-terminal domain (CTD) first directionality [3]. Not only are SF4 helicases well characterized, but *E. coli* is a well-tested model organism, making this an ideal system for investigating replisome mechanics *in vivo*.

DnaB acts as a stable replisome hub that assists in organization and coordination of replication fork activities [4–7]. The N-terminal domain (NTD) of DnaB interacts with the DnaG primase to control priming, and the Tau subunit of the clamp loader complex (CLC) physically and functionally connects the leading and lagging strand polymerases to DnaB. In addition to replisome proteins, DnaB interacts with DnaA and DnaC for replication initiation and loading, Tus for replication termination, and is involved in resolution or bypass of certain replication blocks at the fork [2, 8–15]. Therefore, DnaB’s unwinding activity is likely regulated through multidimensional interactions with several replication enzymes for various replisome related purposes. Indeed, distinct modes of unwinding have been observed. Coupled unwinding and synthesis in an active replisome proceeds rapidly at ~1000 nt/s, but during replication initiation in a complex called the preprimosome and when decoupled from the polymerase during replication, the helicase unwinds at a much slower rate of ~35-100 nt/s [7, 16–18]. The mechanism for this observed kinetic regulation is still being understood.

Biochemical and structural investigations of DnaB have revealed two distinct conformers: a constricted state with a narrow central pore for tight interactions with ssDNA and fast unwinding and a slower, dilated state that allows for translocation over duplex DNA [19–21]. The dilated conformer is used for interactions with DnaC for loading at origins, and with DnaG when laying down RNA primers [19–21]. Investigations into sterically restricted dilated or constricted mutants revealed a direct link between conformation, rate of unwinding, and protein-protein interactions *in vitro*. Studies have also shown that both DNA strands contribute to unwinding, and a mechanism involving steric exclusion and wrapping (SEW) of the excluded ssDNA strand contributes to helicase regulation *in vivo* and *in vitro* [22–24].

Previous investigations of targeted mutations that disrupted helicase regulation led to a proposed dual mechanism, wherein conformational changes in the helicase structure stimulated by protein-protein and protein-DNA interactions acted as a molecular switch to conceal or expose the external residues for engagement with the excluded DNA strand [22]. *In vivo*, these targeted helicase regulation-deficient mutations led to substantial amounts of genomic stress and reduced overall strain growth and efficacy. While helicase regulation is important for genomic stability and damage mitigation, it is not known how dysregulated helicase activity alters the chromosome architecture nor what downstream DNA repair pathways are utilized to maintain a functional genome despite more rapid and uncontrolled unwinding.

RecA binds and cooperatively polymerizes on ssDNA to form filaments for the purposes of homologous recombination (HR) repair of double strand breaks (DSBs) and induction of the bacterial SOS response through allosterically inducing autolytic cleavage of the SOS repressor, LexA [25–30]. While there are rare instances of RecA-independent recombinational repair, bacteria lack the non-homologous end joining (NHEJ) pathway as an HR alternative. RecA filamentation on ssDNA is needed to trigger SOS induction, and RecA-dependent recombination is the only efficient way for *E. coli* to repair DSBs [25, 28, 31]. While DNA damage repair proteins and the translesion synthesis (TLS) polymerase IV (Pol IV) have low-level constitutive expression, activation of the SOS response dramatically upregulates multiple proteins important for rapid DNA repair in the face of severe stress, including a host of DNA damage tolerant (DDT) proteins [32]. For one, the TLS polymerase with the least fidelity, Pol V, is induced and activated as part of the late-stage SOS response [33–35]. While it is unclear at what threshold length RecA filaments trigger SOS induction, functional RecA binding and polymerization is necessary for an efficient response to high levels of stress and DNA damage.

Utilizing exterior SEW mutations in DnaB previously shown to disrupt interactions with the excluded strand and enforce a fast constricted unwinding state are now shown to decouple the replisome and limit leading strand synthesis. These same targeted genomic *dnaB* mutations previously shown to have dysregulated unwinding leading to cellular stress and genomic instability are now combined with an inducible plasmid-based switch to turn off RecA filamentation to investigate the importance of HR and SOS in these strains. RecA activity is important for maintaining fast and efficient growth when helicase regulation is impaired. The high mutational frequency in these *dnaB:mut* strains is dependent on mutagenic repair from SOS-expressed proteins induced by RecA filamentation. Addition of various exogenous DNA damage agents to these strains generally sensitized growth further, and inhibition of RecA filamentation had a neutral or slightly negative impact on survival. A novel <u>P</u>ol I d<u>U</u>TP <u>G</u>ap filling (PLUG) assay was used to label single strand gaps *in situ* and found that dysregulated unwinding leads to increased ssDNA gaps *in vivo*, consistent with a model where helicase regulation encourages functional replisome coupling. In the absence of effective RecA filamentation, these ssDNA gaps are effectively converted to double strand breaks (DSBs) as detected by two separate <u>T</u>erminal d<u>U</u>TP <u>N</u>ick <u>E</u>nd <u>L</u>abeling TUNEL assays. This work makes important advancements in our understanding of the impact of helicase regulation on efficient replisome progression and the dependence on RecA to mitigate consequences from replisome decoupling for cell survival.

## RESULTS

### Constricted DnaB mutants contribute to replisome decoupling

Previously, we have shown that several surface mutations on the DnaB helicase can induce a change in the hexameric structure to a more constricted conformation and dramatically increase unwinding activity [22, 23]. The increase in DNA unwinding activity by these DnaB mutants suggested two possibilities when incorporated in the context of a replisome: 1) either faster unwinding leads to faster synthesis or 2) faster unwinding decouples synthesis and unwinding activities resulting in slower synthesis. To determine which of these possibilities occurs, DnaB mutants were incorporated into a complete *in vitro* bacterial replisome system using a circular TFII DNA substrate (**Fig. S1**) to monitor leading strand synthesis (**Fig. 1**) [36].

**Figure 1.**
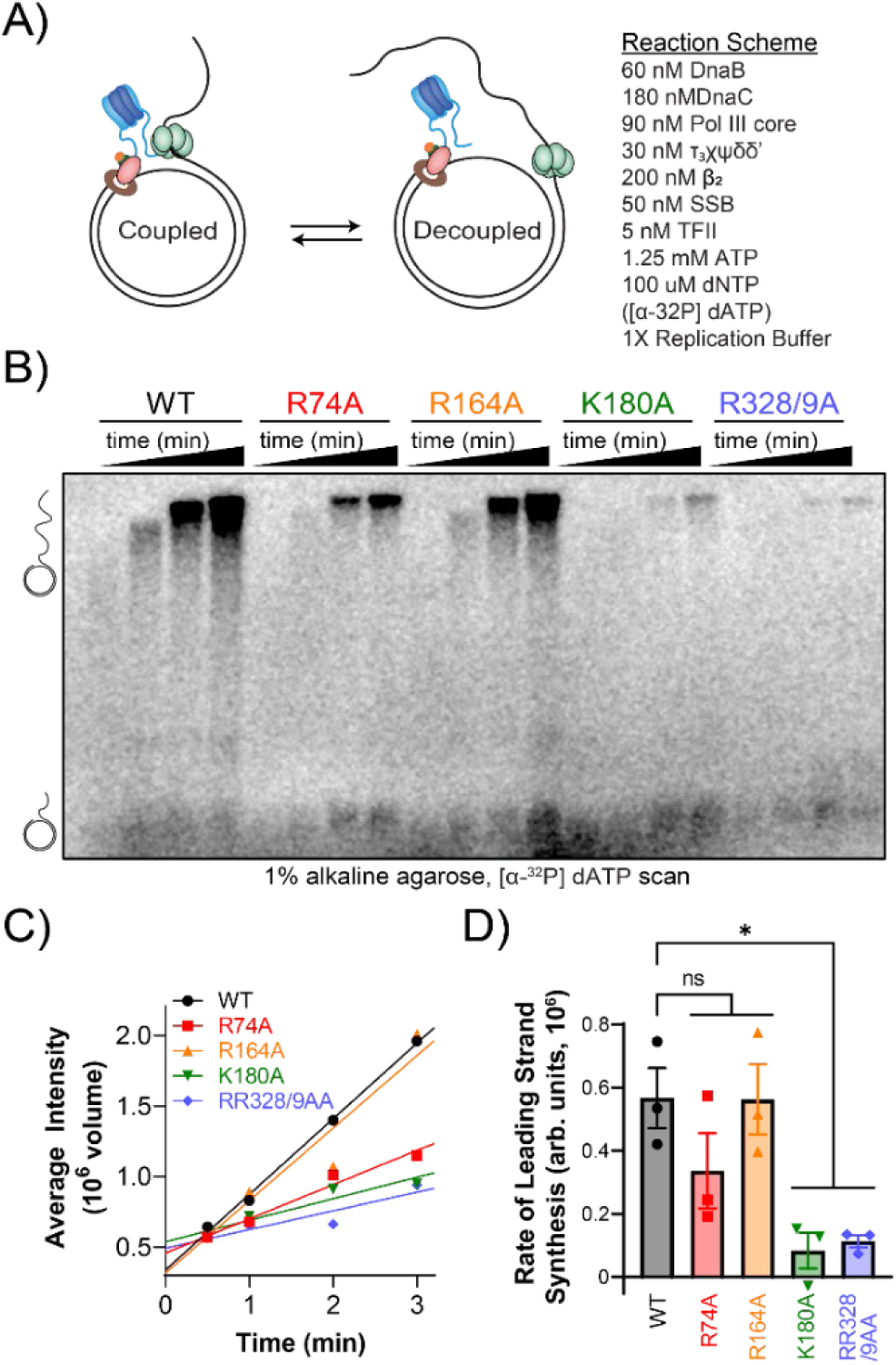
Constricted DnaB mutants decouple the replisome. A) Reaction scheme depicting the replisome components used including TFII substrate (5 nM), 60 nM DnaB_6_, 360 nM DnaC, 30 nM τ_3_χϕδδ’ (CLC), 90 nM Pol III core (αεθ), 200 nM β_2_ in 1X replication buffer at 37 °C for 5 min. The replication reaction was initiated by adding a mixture of 50 nM SSB, 1.25 mM ATP, 125 μM dNTPs, [α-^32^P] dATP. B) ^32^P incorporated leading strand synthesis products with WT or mutant DnaB separated on a 1% alkaline agarose gel and C) quantified at 0.5, 1, 2, 3 mins to calculate the linear rate of synthesis. D) Quantification of the average slope from three independent experiments. Error bars represent the standard error of the mean. Black bars indicate statistically significant differences, where \**P* < 0.05 by a paired two-sided *t*-test.

TFII leading strand reactions utilized DnaC and extended incubation times to allow for maximal loading of all DnaBs prior to initiation [22]. Furthermore, SSB was added in the initiation mix to prevent premature melting of the duplex region. Leading strand synthesis was measured using α-^32^P-dATP incorporation over time. Interestingly, leading strand synthesis, measured by total product abundance, is significantly reduced when fully constricted DnaB mutants (K180A and RR328/9AA) are utilized compared to WT (**Fig. 1B-D**). WT DnaB had a DNA synthesis rate of 5.7 ± 1.0 x10^5^ min^−1^, while moderately constricted mutants, R74A and R164A, had similar rates of 3.4 ± 1.1 x10^5^ min^−1^ and 5.6 x10^5^ ± 1.0 min^−1^, respectively. Alternatively, fully constricted mutants, K180A and RR328/9AA, had significantly reduced rates of synthesis of 0.84 ± 0.06 x10^5^ min^−1^ and 1.1 ± 0.2 x10^5^ min^−1^, respectively, compared to WT.

While the apparent leading strand replication rates were significantly affected with different DnaB mutants, we cannot completely disregard that decreased loading efficiencies or stabilities [22] played a role in these reduced leading strand synthesis rates. The similar DNA synthesis rates observed by both WT and R164A can be a result of the similar DnaB loading efficiencies of the two helicases and effective coupling. The lower synthesis rate of R74A compared to that of R164A or WT can correspond to its slightly reduced (~70%) loading efficiency measured previously [22]. However, the DNA synthesis rates and products were significantly reduced (> 5-fold) for the two fully constricted mutants K180A and RR328/9AA. While some of the reduced DNA synthesis rate of RR328/9AA can be attributed to its inefficiency in loading (~60%), K180A has both a greater loading efficiency (~135%) and a significantly reduced DNA synthesis rate. Therefore, mutations in DnaB (K180A and RR328/9AA) that enforce a persistently constricted state that contribute to decoupled regulation of unwinding and synthesis also create an inefficient replisome.

### RecA filamentation is important for fast, efficient growth when helicase regulation is impaired

As DnaB mutations, K180A and RR328/9AA, decoupled unwinding and synthesis in an *in vitro* assay, and the identical genomic mutations decreased cellular fitness and caused gross chromosomal instabilities [22], we sought to better understand how these *dnaB*:*mut* strains survive in spite of impaired helicase regulation and decoupling by investigating the ssDNA processor RecA. The two constricted DnaB mutant strains *dnaB:K180A* (MSB4) and *dnaB:RR328/9AA* (MSB5), along with the parental strain HME63, were transformed with a plasmid containing an inducible expression system for the dominant mutant allele RecA56 (pEM-RecA56) [37] (see **Table S1** for a full list of strains and plasmids). RecA56 readily incorporates itself into nascent RecA filaments but destabilizes polymerization and prevents filaments from forming on ssDNA [38, 39].

Mass doubling times for all strains were recorded over the course of 20 hours by measuring the *OD_600_* in rich media as performed previously [22]. The tetracycline analog anhydrotetracycline (aTc) was used to induce RecA56 expression in the test strains and showed little to no effect when added at functional doses to control strains, HME63 and the *ΔrecA* strain (JW2669) [40], that lacked the RecA56 plasmid (**Fig. 2A-B**). Adding pEM-RecA56 to HME63 also showed no significant change in growth rate, 0.28 ± 0.01 hr^−1^ and 0.29 ± 0.01 hr^−1^ without and with aTc, respectively (**Fig. 2C**).

**Figure 2:**
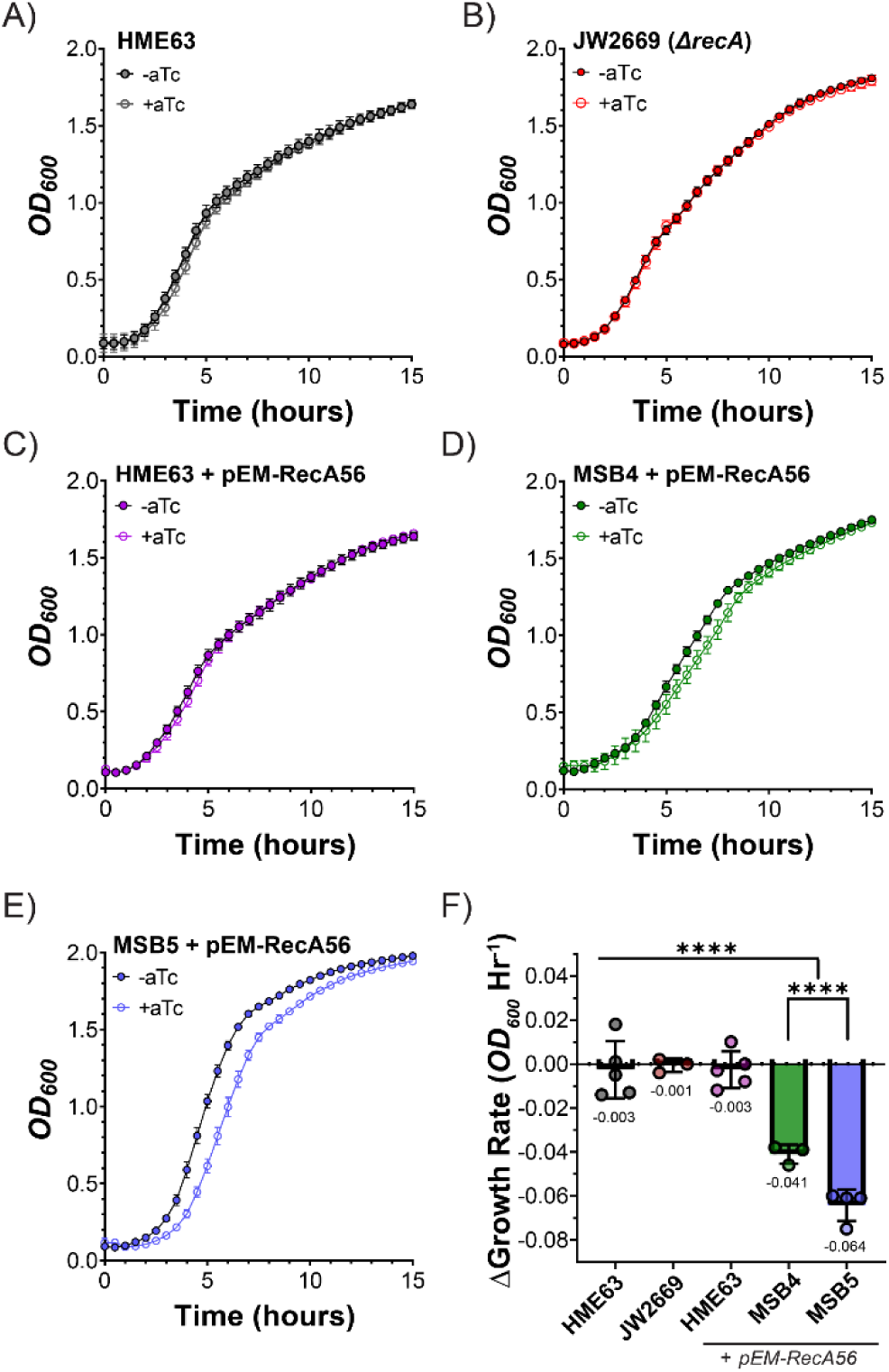
Constricted *dnaB* mutants have a significant reduction in growth rate when RecA filamentation is disrupted. Mean growth curves of each *E. coli* strain A) HME63, B) JW2669 (*ΔrecA*), C) HME63 + pEM-RecA56, D) MSB4 + pEM-RecA56, and E) MSB5 + pEM-RecA56 grown at 32 °C were monitored over a 24-hour period and fitted using Prism 9.5. Individual data is presented with closed (−aTc) or open (+aTc) circles. Error bars represent ± SD and are within the symbols if not visible. E) The log fold change in growth rate from RecA56 expression was measured for each strain and condition by taking the difference of the calculated maximum growth rates ± RecA56 expression. Data is from a minimum of three trials, and individual data is presented with closed circles. Error bars represent ± SD; Black bars indicate statistically significant differences, where p-values are ****< 0.0001, by a paired two-sided *t*-test.

However, RecA56 expression significantly impacted the growth rates of both MSB4 and MSB5 (**Fig. 2D-E**). MSB4 had the slowest growth rate of all the strains, increasing in density at a lower rate than even the RecA knockout, JW2669. During log phase, MSB4 grew at 0.25 ± 0.01 hr^−1^ in the absence of aTc and was reduced by almost 16% to 0.21 ± 0.01 hr^−1^ when RecA56 was expressed with addition of aTc (**Fig. 2D** & **Fig. S2**). Similarly, despite MSB5 demonstrating fast growth with an increase in density of 0.48 ± 0.01 hr^−1^ under normal conditions, expression of RecA56 reduced the growth by ~13% to a rate of 0.42 ± 0.01 hr^−1^ (**Fig. 2E** & **Fig. S2**). To better highlight the different effects of RecA56 expression on these strains, the change in growth rate with aTc added was plotted (**Fig. 2F**). MSB5 had the largest decrease in growth rate, −0.06 ± 0.00 hr^−1^, followed by MSB4 at −0.04 ± 0.00 hr^−1^. However, even though MSB5 had the larger decrease in growth rate compared to the other strains, the growth rate of MSB4 decreased by a greater percentage relative to its unperturbed growth.

### The high mutagenesis of *dnaB* mutant strains is RecA-dependent

Previously, we showed that interfering with helicase regulation led to poor competitiveness in mixed cultures and a dramatic increase in mutational frequencies [22]. From these initial investigations, it was unclear whether the increased mutagenesis was from SOS induction that increased the concentration of low fidelity repair proteins, or some other aspect of helicase dysregulation, perhaps altering the fidelity of Pol III or inducing RecA-independent recombination from excessive ssDNA. To determine if the mutagenesis was RecA dependent, we tested the strains’ abilities to develop resistance to rifampicin with and without expression of the RecA filament interrupter RecA56 (**Fig. 3**). Briefly, cells grown in aTc for RecA56 induction were exposed to rifampicin, and number of resistant colonies (rif^R^) that arose determined the mutational frequencies [41–43]. MSB5 had the greatest number of mutation events under normal conditions, with 8.36 ± 0.45 per million (10^6^) colonies but then drastically dropped to 0.42 ± 0.21 when RecA filamentation was impaired by RecA56 expression (**Fig. 3**, *blue*, solid vs. hashed). MSB4 showed a similar dramatic decrease in mutation frequency when aTc was added, from 4.07 ± 0.26 mutation events per 10^6^ viable cells to 0.73 ± 0.17, indicating a clear reliance on RecA-dependent pathways for mutability (**Fig. 3**, *green*, solid vs. hashed). While the control strain HME63 showed no difference in mutation rate without the RecA56 expression system (**Fig. 3**, *black*, solid vs. hashed), inhibition of RecA by the addition of aTc (*purple*, solid vs. hashed) caused a reduction in mutagenesis from 0.92 ± 0.36 to 0.29 ± 0.13 with aTc. Even in the absence of endogenous stress, RecA allows low level mutagenesis important for environmental adaptation.

**Figure 3:**
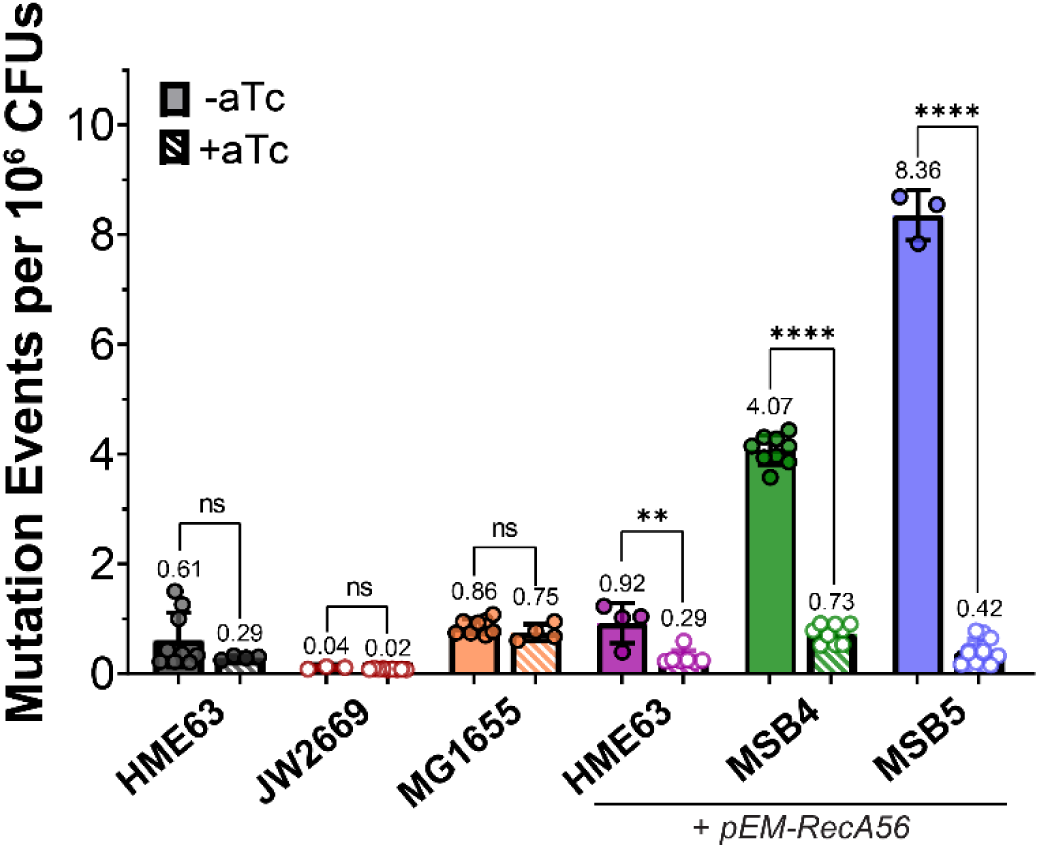
Inhibition of SOS by expression of RecA56 restores basal mutation rates in *dnaB* mutant strains. The mutagenicity of each strain (± pEM-RecA56 and ± aTc, as indicated) was monitored using a rifampicin resistant assay to quantify number of *rif^R^* colonies after 24-hour growth in LB. Average number of resistant colonies per 1 million CFU’s are shown. Data is from at least two trials of three technical replicates each (n ≥ 6) with mean values listed and is presented with closed or open circles indicating functional RecA filamentation of not, respectively. Hashed bars represent addition of aTc to induce RecA56 and inactivate RecA filaments. Error bars represent ± SD; Black bars indicate statistically significant differences, where p-values are **< 0.01 and ****< 0.0001, by a paired two-sided *t*-test.

Interestingly, MSB5 not only had the largest drop in mutational frequency with RecA56 expression but was also reduced by the greatest percentage, dropping by 7.94 events (95%), followed by MSB4 which dropped by 3.34 (82%), and the parental strain dropped by 0.63 mutation events (69%) (**Fig. 3**). Direct comparison of the strains in the absence of aTc showed that the presence of the RecA56 expression system alone did not affect mutation rates, as HME63 ± pEM-RecA56 did not significantly change in mutagenesis (**Fig. S3A**, lanes 1 vs. 4, *solid grey* and *purple*), and the average number of mutations for MSB4 and MSB5 are consistent with previously reported values [22]. Induction of RecA56 to disrupt RecA filamentation resulted in all strains having mutation frequencies less than 1 event per million cells (**Fig. S3B**) showing a clear dependence on RecA-associated pathways for mutation.

### RecA56 expression has a nominal effect on exogenous damage survival for *dnaB:muts* relative to the parental strain

As we determined that the dysregulated helicase strains (MSB4 and MSB5) have high levels of RecA-dependent mutagenesis, we wanted to further explore how they responded to different types of exogenous damage, and whether RecA plays a role in low level genotoxin evasion and survival. We selected three genotoxins to test that elicit different DDT mechanisms: mitomycin C (MMC), which causes large DNA adducts and interstrand crosslinks, methyl methanesulfonate (MMS), which methylates purine bases to cause mispairing, and hydroxyurea (HU) which inhibits ribonucleotide reductase to cause polymerase stalling due to depleted nucleotide pools. Together, these agents provide examples of large blocks to replication that can limit helicase progression, small base changes that are easily bypassed by the helicase but cause polymerase stalling, and an indirect genotoxin that stalls the polymerase without damage, respectively.

Briefly, all cultures were grown overnight in the presence of aTc to ensure strong RecA56 expression in plasmid carrying strains prior to genotoxin exposure. Cultures were serial diluted and spotted (left to right) onto plates containing both aTc and the indicated genotoxin and imaged after 24 hours (**Fig. 4A** & **Fig. S4**). The log-fold survival of MSB4 and MSB5 was significantly impaired relative to the parental strain, HME63, differentially with various genotoxins. Growth of MSB5 was most affected with MMC (**Fig. S4A)**, while growth of MSB4 was severely inhibited with MMS and moderately with both MMC and HU (**Fig. 4B** & **Fig. S4B-C**).

**Figure 4:**
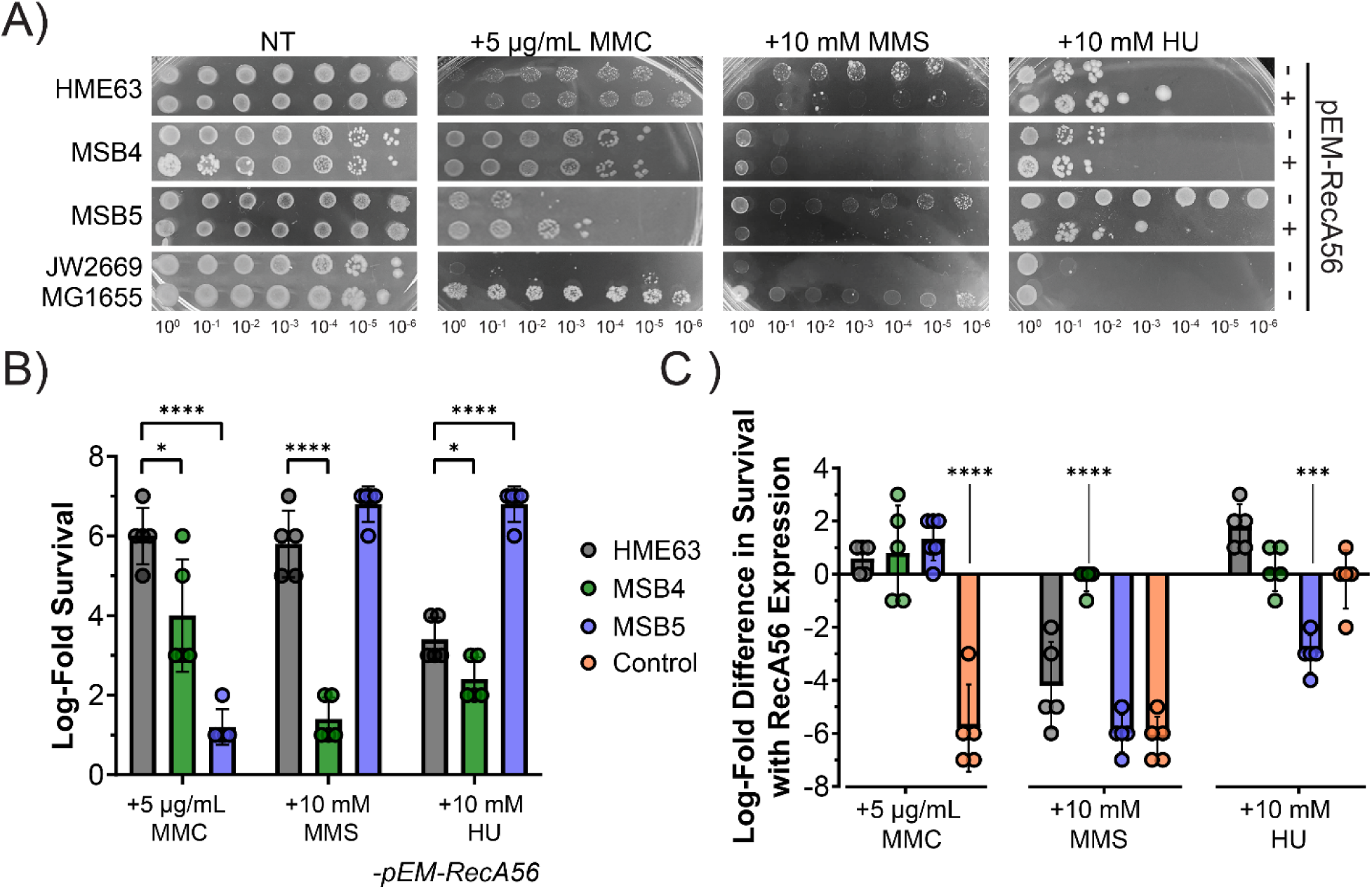
Functional RecA activity is important for low-level genotoxin evasion and survival. A) Representative serial dilution and spotting plates under normal (*far left*) and targeted stress conditions (MMC, MMS, HU) with aTc in the presence or absence of pEM-RecA56 as indicated. B) Quantified log-fold survival in the absence of pEM-RecA56 for each strain and condition. C) Quantified difference in log-fold survival with RecA56 expression for each strain. Control refers to the difference in growth between MG1655 and JW2669. Data was obtained by subtracting log-fold survival in the absence and presence of RecA56. n ≥ 4 trials with individual data points represented as closed circles. Error bars represent ± SD. Black bars indicate statistically significant differences where p-values are *< 0.05, ***< 0.001, and ****< 0.0001 by a paired two-sided *t*-test.

The log-fold difference in survival was plotted in the presence and absence of pEM-Rec56 to compare each strain’s dependence on RecA filamentation for survival when targeted with various genotoxins (**Fig. 4C**). As a control, MG1655 was compared to JW2669 (*ΔrecA*), which shows severe growth inhibition with MMC and MMS. Expression of RecA56 in the presence of MMC caused a slight but nonsignificant increase in log fold survival for both MSB4 and MSB5 (**Fig. S4A**). The largest difference observed was with the addition of MMS, where both HME63 and MSB5 showed significant log fold decreases in survival (−4.2 ± 1.5 & −6.0 ± 0.6, respectively) with RecA56 induction (**Fig. S4B**). Interestingly, MSB5 survived extremely well in the presence of HU compared to either HME63 or MSB4, although inhibition of RecA filamentation by induction of RecA56 reduced the survival equivalent to the parental strain (**Fig. S4C**). Of all the strains tested, MSB4 was the only strain on which RecA56 expression had no significant effect regardless of genotoxin.

### RecA activity is critical for preventing DNA breaks caused by dysregulated unwinding

Because RecA activity was influential for growth and mutagenesis in the *dnaB* mutant strains, and because both MSB4 and MSB5 were sensitive to MMC, we investigated the importance of functional RecA filamentation activity on the frequency of DSBs when DNA unwinding regulation is impaired. MSB4, MSB5, and control strains (HME63 and MG1655), were grown in LB ± aTc until mid-exponential phase, fixed, and permeabilized. <u>T</u>erminal Brd<u>U N</u>ick <u>E</u>nd <u>L</u>abeling (TUNEL) was used to label free 3’-OH ends for detection of both ss- and dsDNA breaks (**Fig. S5A**). Cells were stained with 4′,6-diamidino-2-phenylindole (DAPI, microscopy) or Sytox Green (SG, fluorescence activated cell sorting [FACS]) for nucleotide staining. As a positive controls, HME63 and MG1655 were exposed to MMC for 45 minutes prior to harvesting (**Fig. S6**). Fixed BrdU-labeled cultures were imaged for cellular area and DNA breaks by fluorescence microscopy, with quantification of visual images for MSB5 and HME63, and quantification by FACS for all strains (**Fig. 5B-E**).

**Figure 5:**
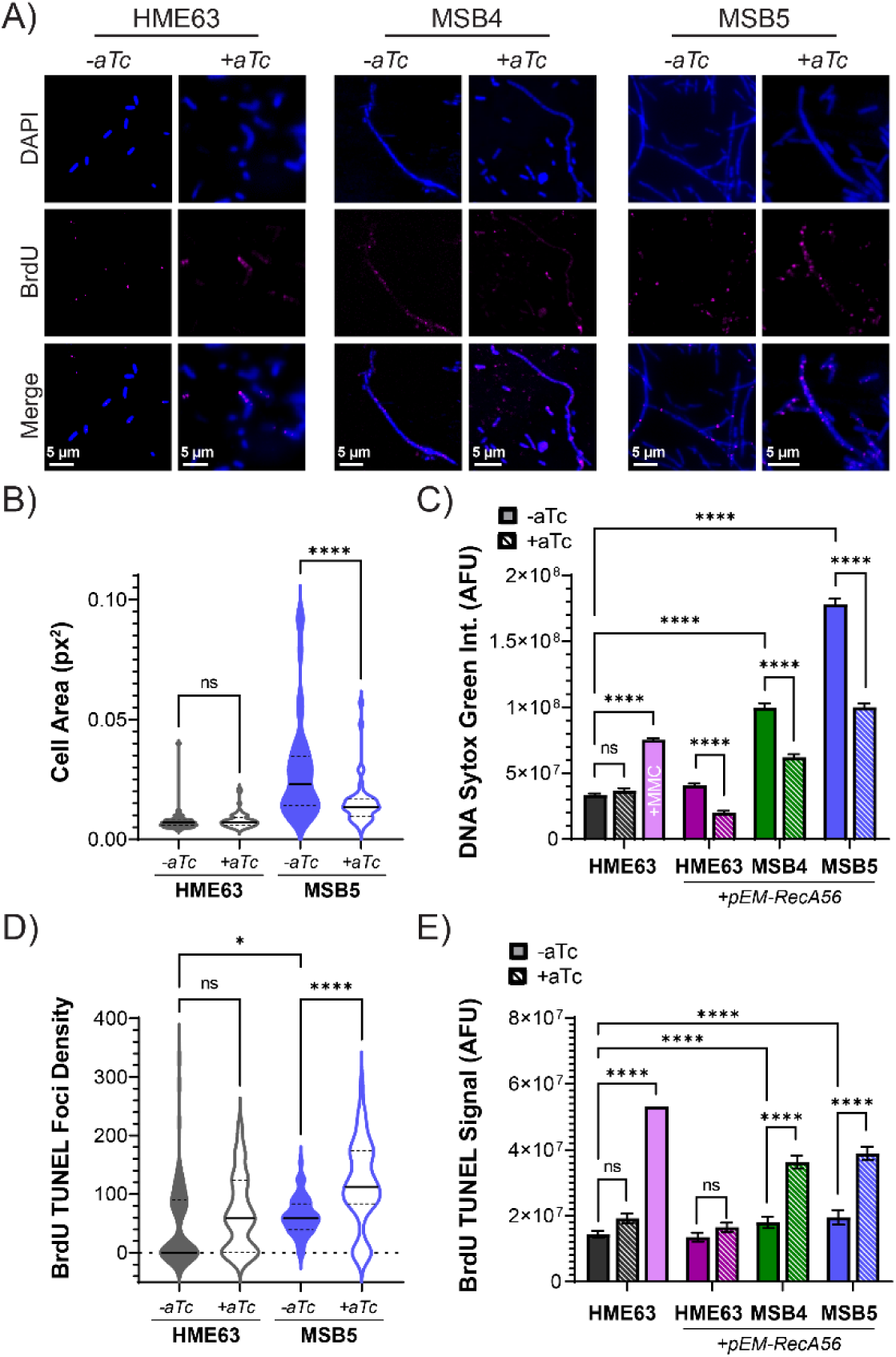
RecA filamentation reduces the amount of DSBs in *dnaB* mutant strains. A) Log phase cells grown in ± aTc for RecA56 induction were stained with DAPI (blue) and BrdU (pink) for *in situ* detection of DNA nicks and breaks by TUNEL. Labeled cells containing the pEM-RecA56 plasmid were imaged by epifluorescence microscopy and images shown are representative of the observed population. Images of control strains exposed to the genotoxin MMC can be found in **Supplemental Figure S6**. The cell area was quantified from B) microscopy for HME63 and MSB5 (px^2^) for more than 50 cells and presented in violin plots, where the solid black bar represents the median and the dashed lines are quartiles in the absence (solid fill) and presence (open fill) of aTc. C) FACS quantifying cellular DNA content using Sytox Green (SG) autofluorescence units (AFU) in the absence (solid fill) and presence (hashed fill) of aTc from n = 10,000 events. Error bars represent ± SD. The intensity of TUNEL BrdU labeling was measured and quantified by D) microscopy as BrdU density to adjust for cell size or by E) flow cytometry (FACS) using the total internal fluorescence for the gated population reported in auto fluorescence units (AFU) from the same exponential growth cultures. Black bars indicate statistically significant differences where p-values are *< 0.05 and ****< 0.0001 by a paired two-sided *t*-test.

HME63 showed mostly normal, short (~2 µm) cells (**Fig. 5A**, *left*) with the occasional filament, that was unchanged with addition of aTc (**Fig. 5B**, *grey*) and contained minimal BrdU foci (**Fig. 5D**, *grey*). However, both MSB4 and MSB5 had a high density of long filamented cells under normal growth conditions, as reported previously (**Fig. 5A**, *middle* and *right*) [22], but these mutant strains appeared to shorten when aTc was added. Interestingly, quantification of MSB5 microscopy images showed a significant ~1.7-fold reduction in cell area when aTc was added (**Fig. 5B**). To more efficiently measure cell size for all strains and conditions, cells were stained with SG and the DNA content was quantified and reported in auto fluorescence units (AFU) by FACS (**Fig. 5C and Fig. S7**). Again, for the control strain, HME63, addition of aTc in the absence of the RecA56 plasmid showed no difference in DNA content per cell; however, addition of MMC significantly increased the SG AFU signal ~2-fold, indicating a high level of cellular stress. MMC is known to induce DSBs, which could increase the DNA content by generating cellular stress from complicated DNA replication processes and inhibited cell division by the SOS protein, SulA, or asymmetrical segregation of Holliday junction entangled chromosomes [44–48]. Interestingly, the trends observed by microscopy quantification for HME63 and MSB5 were confirmed by FACS, and showed a significant 2, 1.6, and 1.8-fold reduction in SG AFU upon aTc addition for HME63, MSB4, and MSB5, respectively.

As a decrease in cellular size was readily observed with addition of aTc, the BrdU TUNEL foci density was quantified for HME63 and MSB5 from microscopy images (**Fig. 5D**). First, in the absence of aTc, there was a significant increase in BrdU staining for MSB5 compared to the control strain, HME63, indicating that even under ideal conditions, DNA breaks are more prevalent in this strain (*grey* vs. *blue, solid fill*). The BrdU TUNEL foci density measured by microscopy was significantly increased with addition of aTc for MSB5, indicating that a reduction in RecA filamentation affords more DNA breaks (*blue, solid* vs. *open fill*). To better quantify the BrdU TUNEL signal in all strains and conditions, FACS was once again utilized (**Fig. 5E**). Consistent with the microscopy quantification, the addition of aTc had no effect on BrdU AFU for the control strain regardless of RecA56 expression, HME63 (*grey or purple, solid vs. hashed*). However, addition of MMC as a positive control for DNA breaks increased the signal >2.7-fold to 5.6 x 10^7^ AFU (*pink*). Both MSB4 and MSB5 had small but significant increases in TUNEL BrdU AFU (1.3 and 1.4-fold, respectively) in the absence of aTc compared to HME63 (*green vs. grey* and *blue vs. grey, solid)*. Interestingly, the AFU signal increased dramatically (~2-fold for both) with addition of aTc to 3.6 x 10^7^ (*green, solid vs. hashed*) and 3.9 x 10^7^ (*blue, solid vs. hashed*), respectively, near that of the +MMC positive control (*pink, solid*).

### *dnaB* helicase mutant MSB4 and MSB5 induce abundant daughter strand gaps

To determine if dysregulated helicase regulation led to the production of excess ssDNA gaps *in vivo* stimulating the protective activities of RecA, we utilized Klenow, an exonuclease deficient variant of Pol I, in a <u>P</u>ol I d<u>U</u>TP <u>G</u>ap filling (PLUG) assay (**Fig. S5B**) [49–51]. Briefly, the same cells that were probed with TUNEL (**Fig. 5**, *above*) were split and incubated at room temperature for 30 minutes with Klenow and a nucleotide mix containing BrdU to label any ssDNA gaps extended from a 3’-OH primer. To limit variables, PLUG-treated cells were analyzed by microscopy and FACS identically to TUNEL samples. The amount of PLUG signal detected in the control strains, HME63, MG1655, and JW21669, was minimal and did not appear to change with addition of aTc (**Fig. 6A** & **Fig. S6**). However, the addition of MMC to HME63 caused a visual increase in PLUG signal, indicating a substantial amount of genomic stress (**Fig. S6**). Qualitatively, PLUG foci were generally brighter than TUNEL, likely due to multiple BrdU nucleotides being incorporated into a single ssDNA gap. Similar to TUNEL, PLUG foci and cell area were measured and used to calculate and compare PLUG density for HME63 and MSB5 (**Fig. 6B**). The PLUG foci density did not significantly change for HME63 or MSB5 when aTc was added (*solid vs. open fill, blue or grey*), possibly from the limited number of quantifiable cells, as the median value for MSB5 appeared greater than for HME63.

**Figure 6:**
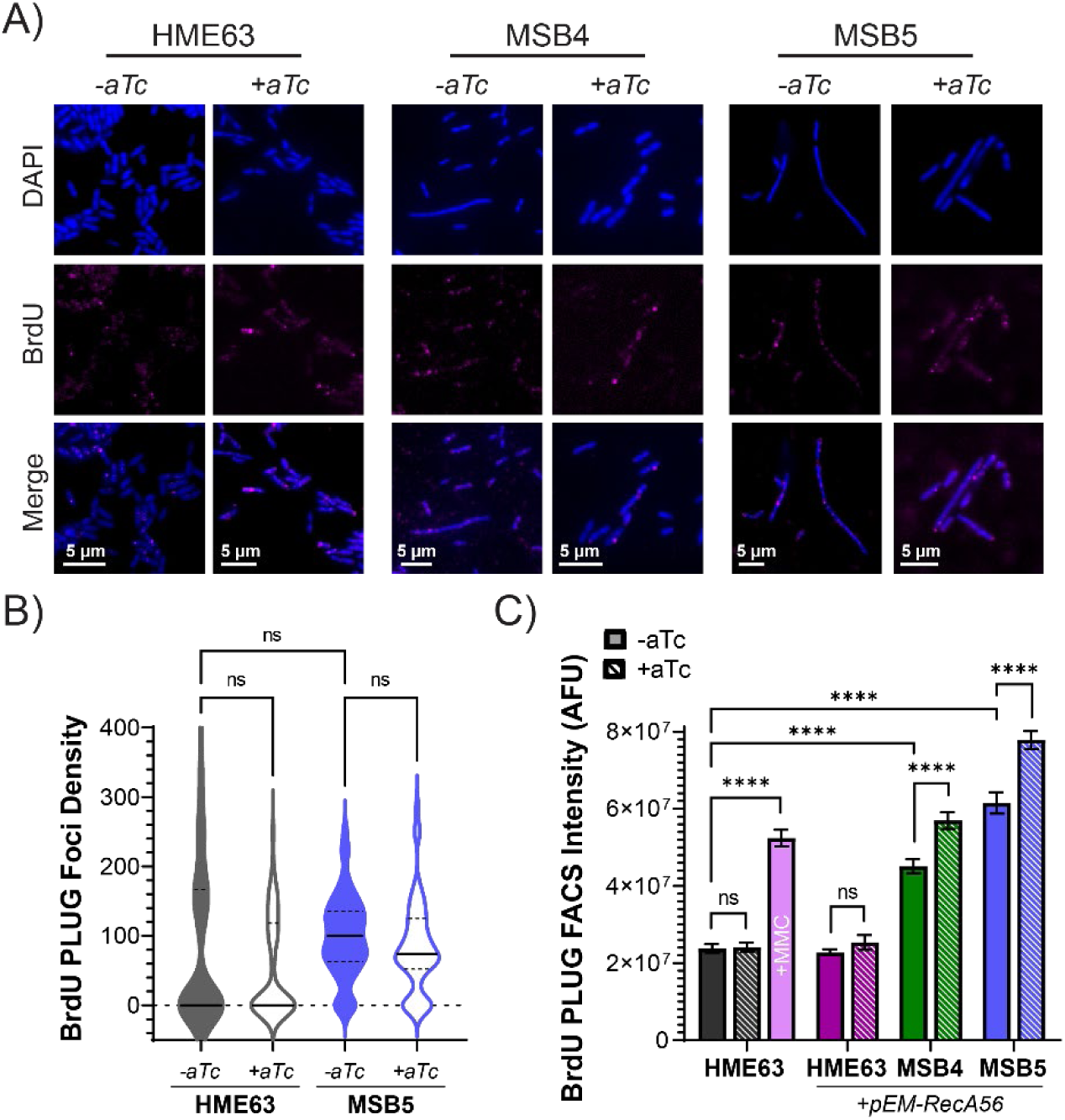
Dysregulated helicase activity induces significant ssDNA gaps. A) Log phase cells grown in ± aTc for RecA56 induction were stained with DAPI (blue) and BrdU (pink) for *in situ* detection of DNA nicks and breaks by PLUG. Labeled cells containing the pEM-RecA56 plasmid were imaged by epifluorescence microscopy and images shown are representative of the observed population. Images of control strains exposed to the genotoxin MMC can be found in **Supplemental Figure S6**. The intensity of PLUG BrdU labeling was measured and quantified by B) microscopy as BrdU density to adjust for cell size, where the black bar represents the median in the absence (solid fill) and presence (open fill) of aTc or by C) flow cytometry (FACS) using the total internal fluorescence reported for the gated population in auto fluorescence units (AFU) in the absence (solid fill) and presence (hashed fill) of aTc from n = 10,000 events. Error bars represent ± SD. Black bars indicate statistically significant differences where p-values are *< 0.05 and ****< 0.0001 by a paired two-sided *t*-test.

To better quantify PLUG signal intensity across multiple strains and conditions for larger sample sizes, the BrdU PLUG AFU signal intensity was quantified per cell by FACS (**Fig. 6C**). First, the PLUG intensity measured for MSB4 (4.5 x 10^7^ AFU) and MSB5 (6.2 x 10^7^ AFU) was 1.9 and 2.6-fold greater than for HME63 (2.4 x 10^7^ AFU) respectively, indicating that an excessive number of single strand gaps are created when the helicase is dysregulated. The single strand gaps measured by PLUG AFU (**Fig. 6C**) are ~ 2-fold more intense in all cases compared to DNA breaks measured by TUNEL AFU (**Fig. 5E**) when DnaB is dysregulated. Although there was no difference in HME63 ± aTc (*grey or purple, solid vs. hashed*), the addition of aTc to MSB4 or MSB5 significantly increased the PLUG AFU further 1.3-fold for both (5.7 and 7.8 x 10^7^ AFU, respectively, *green or blue, solid vs. hashed*). The increased change in AFU for PLUG (**Fig. 6C**) is of less magnitude than for TUNEL (**Fig. 5E**) when RecA filamentation is disrupted. This result combined with the decrease in cellular area with RecA disruption (**Fig. 5C**) allows us to conclude that dysregulation of helicase unwinding induces long stretches of ssDNA that are protected and managed by RecA. However, when RecA filamentation is disrupted, these ssDNA gaps are converted to DNA breaks that are likely processed through a separate DDT pathway that allows for cell division.

## DISCUSSION

In a previous study, we described the first *in vivo* investigation of genomic *dnaB* SEW-disrupting mutations, reporting that MSB4 (*dnaB:K180A*) and MSB5 (*dnaB:RR328/9AA*) stabilized a constricted DnaB conformer and had significant negative impact on replication fidelity and cellular efficacy [22]. Both strains demonstrated a filamented cellular stress phenotype and had high incidence of mutations, implying a dependence on RecA for survival. Here, these constricted and fast unwinding mutants of DnaB (K180A and RR328/9AA) both drastically reduced productive leading strand synthesis products, implying that coupling between unwinding and synthesis must be maintained for a productive replisome. In order to better understand helicase regulation *in vivo* and the consequences of dysregulation, we investigated the role of RecA filamentation on strain survival and genomic stability using an inducible system to inhibit RecA activity through the addition of aTc and expression of RecA56. The high mutagenicity of *dnaB* mutant strains [22] was found to be dependent on RecA polymerization and is therefore the result of increased DDT from SOS protein induction, including mutagenic Pol V. High levels of endogenous damage in *dnaB* mutant strains that presented as DNA nicks and gaps were tolerated by RecA filamentation for growth and efficacy, albeit with a cellular stress and filamentation phenotype. When exposed to exogenous damage, dysregulated unwinding in these strains typically increased genotoxin toxicity, but RecA activity had an overall neutral or positive effect on survival. Interestingly, dysregulated helicase activity on its own produced excess ssDNA *in vivo*, and RecA-mediated HR is partially responsible for the cell filamentation and asymmetric chromosome segregation observed in these strains [22]. Importantly, RecA filamentation is utilized to modulate ssDNA gaps created by dysregulated helicase activity, and inhibition of RecA filamentation in these strains converts ssDNA gaps to breaks allowing for alternative DDT mechanisms and cellular division (**Fig. 7**).

**Figure 7:**
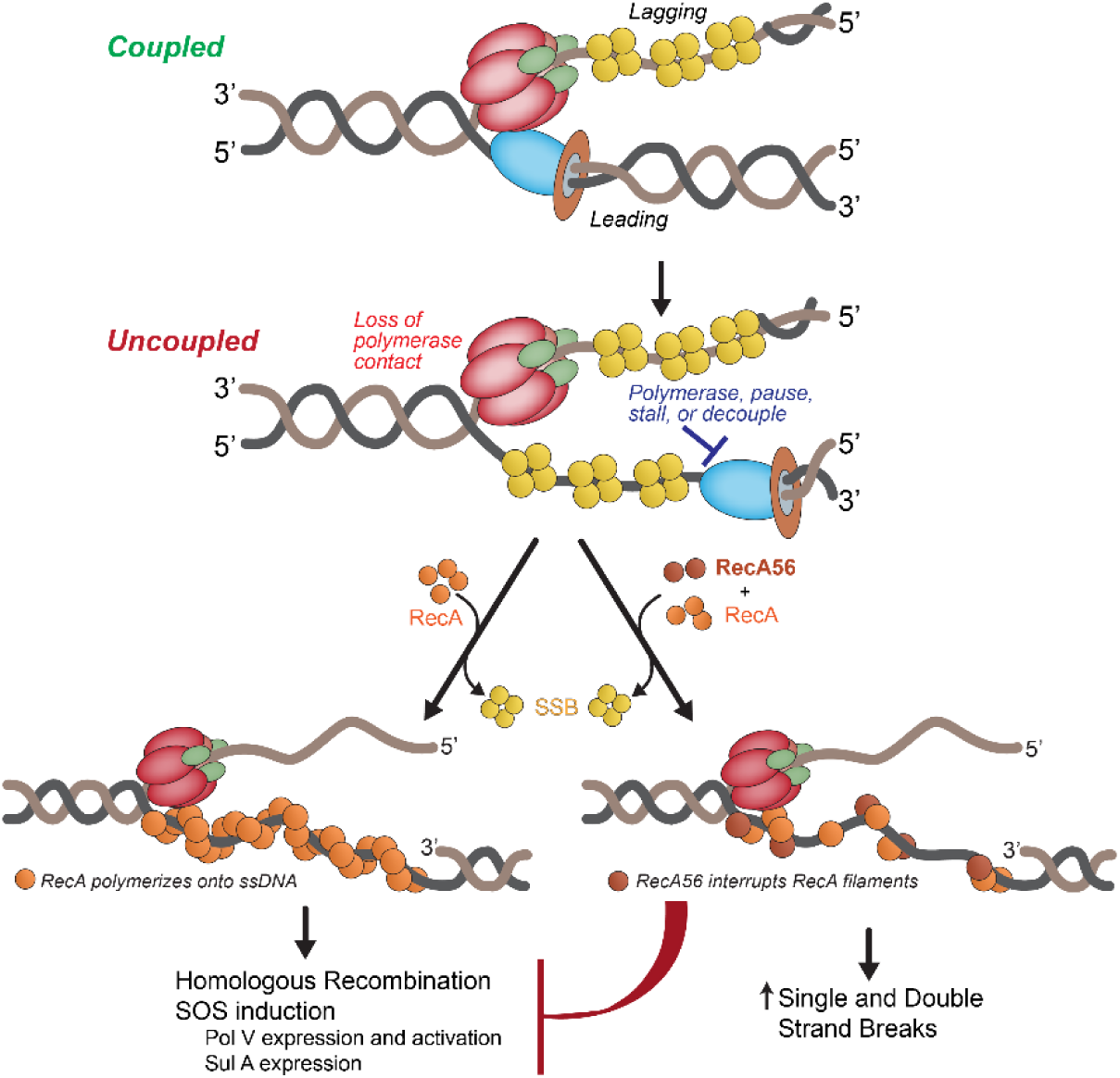
Helicase polymerase uncoupling induces ssDNA gaps that stimulate RecA filamentation to protect and repair the genome. Uncoupling of unwinding and synthesis from helicase dysregulation or other persistent blocks to replication lead to an increased frequency of ssDNA gaps that are modulated by RecA filamentation and downstream SOS and DDT processes. However, when RecA56 is induced, ssDNA gaps are unprotected, become labile, and are converted to more single and double strand breaks.

Although MSB5 had a significantly higher mutation frequency than MSB4, both had increased RecA-dependent mutagenesis from increased SOS induction relative to the parental strain, HME63. When RecA56 was induced to impair RecA filament formation, both *dnaB* mutants demonstrated fewer than 1 in 10^6^ mutation events, equivalent to parental strains, revealing that the increased mutation frequency of *dnaB* mutant strains can be almost entirely attributed to SOS induction through productive RecA filamentation. Exposure with targeted exogenous damage showed that the high mutational frequencies of the dysregulated helicase strains did not functionally contribute to genotoxin evasion or survival. However, MSB5 with its greater SOS induction [22] was more effected by MMC than MSB4 or HME63. Generally, RecA56 expression had a neutral or negative impact on survival for all strains, indicating a RecA polymerization-dependent mitigation of damage was independent of DDT, SOS protein induction, and repair. Interestingly, MSB5 was resistant to HU treatment but was sensitized when RecA filamentation was inhibited by RecA56. HU functions to stall polymerization through reduction in available dNTPs, and as TLS Pol V has a lower nucleotide *K_d_*, the upregulated SOS protein expression in MSB5 [22] allows for more consistent TLS polymerase fork rescue and extension. Polymerase exchange at the fork with higher concentrations of mutagenic TLS Pols likely allows for frequent and continuous replication by Pol V within large stretches of ssDNA gaps generated by MSB5 [52–55]. Since dysregulated unwinding stimulates RecA polymerization on ssDNA gaps causing induction of SOS, we were interested in whether the growth of *dnaB* mutant strains during RecA inhibition impact strain growth. RecA56 expression significantly reduced the growth rate of *dnaB* mutant strains. Therefore, dysregulated *dnaB* mutants depend on RecA activity to mitigate the impact of rapid unwinding *in vivo*.

Previously we reported that MSB4 and MSB5 had increased chromosome complexities and complementary increases in the *ori:ter* ratios, suggesting that fork stalling from decoupling was impacting chromosome segregation [22]. Quantification of the large increase in DNA breaks and total DNA intensity in *dnaB* mutant strains relative to the parental shows that helicase-affected DNA damage and fork stalling forces cell filamentation and asymmetrical separation of chromosomes. With RecA, the majority of measured TUNEL foci correlate directly with DSBs and are therefore substrates for HR repair, which are known to create temporarily entangled chromosomes through RecA-mediated strand invasion [47, 56]. While inhibition of RecA polymerization reduced the DNA content and filamentation of *dnaB* mutant cells, they still maintained a higher DNA content than the control strains, indicating that not all chromosome entanglement and asymmetric segregation is from active Holliday junctions or that not all Holliday junctions are fully resolved. Stretches of ssDNA induced from decoupling are also converted to DSBs in the absence of productive RecA filamentation. ssDNA secondary structures, DNA adducts, and other lesions are likely also involved, delaying fork progression to the terminus and sister chromosome separation.

This dependence on RecA activity to maintain fast and efficient growth when helicase activity is dysregulated or impeded indicates that measurably more endogenous damage is constantly challenging faithful and efficient replisome progression. Both DnaB K180A and RR/328/9AA helicase mutants failed to produce long leading strand products *in vitro*, suggesting that these mutants are causing significant decoupling of unwinding and synthesis, comprising the replisome. This observed decoupling *in vitro* is further supported by excessive DNA breaks and even more ssDNA gaps detected endogenously by TUNEL and PLUG when helicase regulation is disrupted *in vivo* (*i.e.* MSB4 and MSB5). Direct comparison of TUNEL and PLUG between MSB4 and MSB5 shows that despite the high incidence of DNA breaks, DNA gaps are the primary product of dysregulated unwinding. When faced with rapid ssDNA generation, RecA-associated repair is critical for cell survival, efficient replication, and damage mitigation in *E. coli* [57]. Even though complex and diverse fork interactions maintain a dynamic replisome, helicase regulation is critical for optimal replication and genome duplication. Targeted regulation-deficient helicase mutants induce replisome decoupling *in vitro* and *in vivo*, but are biologically mitigated by RecA stabilization and recombination.

## MATERIALS AND METHODS

### Materials

ATP was from Invitrogen (Carlsbad, CA). α-^32^P-dATP was from Perkin Elmer (Waltham, MA). DNA substrates were HPLC or gel purified from Sigma-Aldrich (St. Louis, MO) or IDT (Coralville, IA). Mitomycin C (MMC) (Fisher Bioreagents, Waltham, MA) and Hydroxyurea (HU) (Acros Organics, Waltham, MS) were dissolved in DMSO; anhydrotetracycline (aTc) (Thermo Fisher Scientific, Waltham, MA) was dissolved in ethanol; Methyl methanesulfonate (MMS) (Alpha Aesar, Haverhill, MA) was dissolved in ultrapure water prior to use. Rifampicin (rif) (Fisher Bioreagents, Waltham, MA) was dissolved in DMSO. Sytox Green (SG) was from Invitrogen (Carlsbad, CA). All other materials were from commercial sources and were analytical grade or better.

### TFII Substrate Preparation

The tailed form II (TFII) substrate was prepared using the pSCW01 plasmid as described previously [36]. Briefly, pSCW01 was isolated from cell pellets using ZymoPURE plasmid midiprep kit (Irvine, CA). 100 μg of pSCW01 plasmid was treated with 1.5 units/μg of site specific endonuclease *Nt.BstNBI* (New England Biolabs, Ipswitch, MA) and 100X molar excess of displacer oligonucleotides (DNA197-199, sequences complementary to the single stranded DNA fragments created by *Nt.BstNBI* enzyme) in 1X NEB 3.1 buffer at 55 °C for 4 hours to create a gapped plasmid. The *Nt.BstNBI* enzyme was heat inactivated at 80 °C for 20 mins according to the manufacturer’s instructions. In the same reaction tube, the nicked DNA fragments were annealed with the displacer oligos in a thermal cycler at a cooling rate of 1 °C/min until the reaction reached 12 °C. To extract the gapped plasmid, the displacer oligos were removed using PEG purification where an equal volume of 2X PEG solution (26 % [w/v] PEG-8000, 20 mM MgCl_2_) was added to the reaction mixture, centrifuged at 21,000 x g for 1 hr at 6 °C and the supernatant was discarded. The pellet containing the gapped plasmid was washed with 70 % (v/v) ice cold ethanol and centrifuged at 21,000 x g for 30 min at 6 °C. The supernatant was discarded, air dried, and the resulting pellet was resuspended in 80 μl of Milli-Q water prewarmed to 65 °C. The gapped plasmid was then annealed to 3X molar excess of fork oligonucleotide, DNA200 in thermal cycler in 1X Cutsmart buffer at 50 °C for 10 min, followed by cooling to 16 °C at a 1 °C/min cooling rate. Next, ligation was performed by the addition of 62.5 units of T4 DNA ligase per μg of DNA substrate supplemented with 8 mM ATP and 1 mM DTT and incubated at 16 °C for 18 hrs. The ligase was heat inactivated at 65 °C for 20 min according to the manufacturer’s instructions. The TF II substrate was purified as before by the addition of an equal volume of 2X PEG solution, followed by washing with 70 % ethanol. The resulting pellet was resuspended in 1X TE buffer.

### TFII Rolling Circle Assay

The *E. coli* DnaB helicase wild-type and mutants (R74A, R164A, K180A, R328/329A), DnaC loader [22], Pol III core [58], ß-clamp [59], τ_3_ clamp-loader-complex (τ_3_-CLC) [60], and SSB [61] were expressed and purified essentially as previously described. The TFII substrate (5 nM) was incubated with 60 nM DnaB_6_, 360 nM DnaC, 30 nM τ_3_χϕδδ’(τ_3_-CLC), 90 nM Pol III core (αεθ), 200 nM β_2_ in 1X replication buffer at 37 °C for 5 min. The replication reaction was initiated by adding a mixture of 50 nM SSB, 1.25 mM ATP, 125 μM dNTPs, α-^32^P-dATP in 1X replication buffer (50 mM HEPES [pH 7.9], 12 mM Mg(OAc)_2_, 0.1 mg/mL BSA, 10 mM DTT) prewarmed to 37 °C. The replication reaction was stopped by mixing with an equal volume of quench buffer (200 mM EDTA, 300 mM NaOH, 18 % Ficoll [w/v], 0.15 % [w/v] Bromocresol green, 0.25% [w/v] xylene cyanol) at different time points. The reaction products were electrophoresed using 1% alkaline agarose gels at 35V until the dye front ran two thirds of the gel. The gels were dried, exposed overnight, and imaged on a Typhoon FLA 9000 phosphorimager (Cytiva, Marlborough, MA). The bands were quantified using ImageQuant software (v.10.1).

### RecA56 Vector Engineering

To create a plasmid-based RecA56 expression system, the plasmid pEM-Cas9HF1-indRecA56 (Addgene ID: 102294) designed for the co-inducible expression of Cas9 and RecA56 [37] was modified using primers *Cas9HF1delta F* and *R* (**Supplemental Table S2**) to create pEM-indRecA56 (pEM-RecA56). These primers remove Cas9, keeping the *tet* promoter and *recA56* genes intact, allowing for inducible expression solely of the *recA56* gene. This modified plasmid was transformed into Top10 cells and screened by restriction digest and DNA gel electrophoresis, before being electroporated into HME63, MSB4, and MSB5.

### Growth Curves

Growth curves were recorded by diluting overnight clonal cultures 1:100 (*OD_600_* ~ 0.01) in LB and aliquoting 250 μL into a round bottom clear 96-well plate (Corning) with or without the addition of 100 ng/mL anhydrotetracycline (aTc). The cultures were incubated at 32 °C (to prevent λ-Red induction in these strains) with aeration at 225 RPM and the *OD_600_* was recorded at 30-minute intervals using a Tecan Spark microplate reader equipped with Spark Control Magellan software (Tecan, Männedorf, Switzerland). Peak growth rate was determined from the maximal slope using Spark Control Magellan, and data was processed and plotted using Prism 9.5 (GraphPad, San Diego CA).

### Strain Mutagenesis Assay

To determine the mutation frequency of each strain, a rifampicin resistant (*rif^R^*) assay was performed as previously described [62], with the modification of using rich media [22]. To test the mutagenesis rate with and without RecA activity impairment, fresh overnight cultures were diluted 1:100 in LB media with or without the addition of 100 ng/L aTc (Thermo Fisher Scientific, Waltham, MA). Cultures were grown at 32 °C with aeration until *OD_600_* ~ 0.4, then again diluted 1:100 in fresh LB (± aTc) and allowed to grow for 24 hours. 100 µL aliquots were spread onto plates containing 50 µg/mL rifampicin (ThermoFisher, Waltham, MA) (*rif^+^*). Identical aliquots were diluted and plated onto regular LB (*rif^−^*) plates to calculate colony-forming units (CFUs). Plates were incubated at 32 °C until colonies appeared or 48 hours was reached. Mutation frequency was calculated as the ratio of mutants to total CFUs as follows:

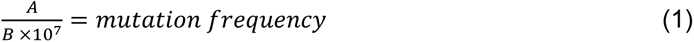

where *A* is the number of mutant CFUs (colonies on the *rif^+^* plate), *B* is the number of total CFUs (colonies on the *rif^−^*plate), and *10^7^*is the dilution factor for *B*.

### Genotoxin survival assays

For cell survival assays, fresh clonal cultures from glycerol stocks were grown overnight at 32 °C in LB medium either with or without 100 ng/mL aTc for induction of RecA56. Cultures were normalized by *OD_600_*, then serially diluted and spotted in 2 μL aliquots onto LB-agar plates containing the indicated concentration of genotoxin (MMC, HU, MMS) and ± 100 ng/mL aTc. Plates were incubated at 23 °C for approximately 1 hour, then transferred to a 32 °C incubator for 24-48 hours before imaging and data collection. All stock solutions were made at 1000X prior to mixing with cooled LB agar. Plate images were taken on the benchtop using a handheld camera; data was processed using Excel and plotted using Prism 9.5 (GraphPad, San Diego CA).

### Fluorescent detection of DNA nicks and gaps *in situ*

Exponential growth cultures were obtained by diluting overnight cultures 1:100 in LB ± 100 ng/mL aTc for RecA56 induction and grown overnight at 32 °C. Overnight cultures were again diluted 1:100 in 50 mL LB ± 100 ng/mL aTc and grown with aeration at 32 °C until OD ~ 0.5. Cells were harvested by pelleting and washing with PBS, then fixed in 1 mL of ice cold formaldehyde solution (4% paraformaldehyde in 1X PBS) for 20 minutes at room temperature, pelleted, and washed with PBS as described [63]. For internal controls, exponential growth cultures of the parental strain HME63, the K12 strain MG1655, and the Keio collection *ΔrecA* strain JW2669 [40, 64] were exposed to 1 µg/mL MMC for 60 minutes prior to harvesting. After fixation, cells were permeabilized by rapid resuspension in 77% ice cold ethanol, then pelleted, washed, resuspended in a PBS storage solution containing 0.1% PFA, and kept at 4 °C.

To visualize DNA breaks, <u>T</u>erminal d<u>U</u>TP <u>N</u>ick <u>E</u>nd <u>L</u>abeling (TUNEL) was utilized to label free 3’ DNA ends [51]. For direct comparison of results, half of each fixed sample was processed for TUNEL analysis, and the remaining half was reserved for single strand gap detection (below). dUTP was added to DNA ends by pelleting stored cells and resuspending in 100 μL of elongation buffer (1X terminal deoxytransferase [TdT] buffer, 2.5 mM CoCl_2_, 0.3 mM 5-bromo-2’-deoxyuridine [BrdU] [Invitrogen, Waltham, MA]), then adding 5 U of TdT (Thermo-Fisher, Waltham, MA), and incubating at 37 °C for 60 min. After elongation, cells were pelleted, washed with PBS, and then resuspended in blocking solution (4% BSA in 1X TBST) for 30 minutes at room temperature. To fluorescently label BrdU labelled ends, blocked cells were pelleted and resuspended in 100 µL of primary antibody solution (1:100 mouse-α-BrdU [BD Bioscience, Franklin Lakes, NJ] in TBST with 4% BSA) for 60 minutes at room temperature. Afterwards, cells were pelleted and washed three times with blocking solution, then resuspended in 100 µL of secondary antibody (1:500 α-mouse IgG-Alexa647 [ThermoFisher, Waltham, MA] in TBST with 4% BSA), and incubated in the dark for 60 minutes at room temperature. Cells were pelleted again, washed with TBST, washed twice with PBS, then resuspended in the PBS storage solution and kept at 4 °C for analysis.

To visualize single stranded DNA, an exonuclease deficient Klenow variant of Pol I (New England Biolabs, Ipswich, MA) was used to extend DNA ends across daughter strand gaps [49, 50] in a <u>P</u>ol I d<u>U</u>TP <u>G</u>ap-filling (PLUG) assay. The reserved half of the stored fixed cells were pelleted and resuspended in 100 μL Klenow extension buffer (1X NEB buffer 2, 10 µM each of dATP, dCTP, dGTP, and BrdU [ThermoFisher]). To initiate the elongation, 5 U of Klenow was added, and the reaction was incubated at room temperature for 40 minutes. After elongation, cells were pelleted, washed thoroughly with PBS, then resuspended in blocking solution (4% BSA in 1X TBST) for 30 minutes at room temperature. Incorporated BrdU nucleotides were labeled as described above for the TUNEL assay. After secondary antibody incubation, cells were washed with TBST, then resuspended in PBS storage solution and kept at 4 °C for analysis by epifluorescence microscopy and flow cytometry.

### Microscopy

All microscopy images were obtained using an Olympus Brightfield Microscope IX-81 (Olympus Corp., Center Valley, PA) using a 60x objective with oil immersion. 2 µL of fixed samples from TUNEL and Klenow extension assays were spotted onto a microscope slide and allowed to dry. DAPI (ThermoFisher, Waltham, MA) was added to mounting media (2.5% DABCO, 90% glycerol, 7.5% PBS) to create a dual staining and mounting solution. 3 µL of this solution was added to cover the fixed cells, then immediately topped with a coverslip and sealed with clear polish. Slides were stored at 4 °C in the dark overnight prior to imaging.

The DAPI stained images were used to set masks for individual cells with manual adjustments made for overlapping cells or those not identified using Image J (v.153) [65]. The area embedded by the masks was utilized to give cell area. Alexa647 foci from TUNEL or PLUG assays were quantified from pixel maxima within the mask to give the foci per cell. Foci per cell was divided by cell area to obtain foci density. This data was analyzed by excel and plotted using Prism 9.5 (GraphPad, San Diego CA).

### Flow Cytometry

To quantify cell size and BrdU intensity, TUNEL and PLUG treated and Alexa647 antibody-stained cultures were pelleted, resuspended in 1 mL sheath fluid (sterile 1X PBS) with 1.5 µM Sytox Green, and incubated in the dark for 30 minutes. Samples were diluted with additional sheath fluid based on cell density and instrument parameters before being analyzed by FACSverse (BD Biosciences). HME63, MG1655, and JW2669 exposed to 1 µg/mL MMC prior to harvesting were used as a positive controls; untreated cells stained with Sytox Green and unstained fluorescently labeled BrdU cells were used as signal controls [51].

Using FloJo software (v10.8.1, BD biosciences), cells were gated for single cells based on a forward scatter height vs. area scatter plot (see **Fig. S7A**) and universal gates for ± BrdU and ± SG subpopulations were set based on allophycocyanin (APC) and fluorescein isothiocyanate (FITC) controls, respectively (**Fig. S7, B-E**). To calculate BrdU intensity for TUNEL and PLUG, FACS histograms of each sample were exported from FloJo as a series of individual values. Each sample’s total fluorescence of +BrdU and +SG gates were independently calculated using Equation 2:

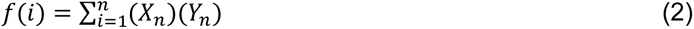

where *i* is the bin number, *X* is the AFU of that bin, *Y* is the cell count of that bin, and *n* is the total number of bins associated with the sample. Total fluorescence values were calculated in Excel and data was plotted using Prism 9.5 (GraphPad, San Diego CA).

## Acknowledgements

Special thanks to Ben Van Houten (U. Pittsburgh) for providing us with pSCW01 and to Charles McHenry (U. Colorado) for providing us with initial *E. coli* replisome proteins, plasmids, antibodies, and cell stocks. We thank all members of the Trakselis laboratory for productive conversations and insight. We acknowledge the Baylor Molecular Bioscience Center (MBC) and the Center for Microscopy and Imaging (CMI) for providing instrumentation and resources aiding this project. This work was funded by the NSF MCB (NSF 2105167 to M.A.T.) and supported by Baylor University.

## Author Contributions

MSB: conceptualization, formal analysis, investigation, validation, visualization, project administration, data curation, methodology, writing—original draft, writing—review and editing; HMP: formal analysis, investigation, methodology; MUW: investigation and formal analysis; JEM: investigation; MAT: conceptualization, formal analysis, investigation, visualization, methodology, supervision, funding acquisition, project administration, resources, writing— original draft, writing— review and editing

